# Silmitasertib, an FDA-designated orphan CK2 Inhibitor, ameliorates neuropathology and motor dysfunction in a Huntington’s disease mouse model

**DOI:** 10.1101/2025.11.19.689275

**Authors:** Ross J. Pelzel, Miaya A. Herbst, Nicholas B. Rozema, Melissa A. Solem, Rocio Gomez-Pastor

**Author notes:** Corresponding author: Rocio Gomez-Pastor, Department of Neuroscience, University of Minnesota, School of Medicine, 2101 6th St. SE, Minneapolis, MN 55544, USA.

## Abstract

Huntington’s disease (HD) is a devastating autosomal dominant neurodegenerative disease that manifests with progressive motor, cognitive, and psychological impairments. HD is caused by a polyQ (CAG) repeat expansion in the huntingtin (*HTT*) gene, leading to the misfolding and aggregation of mutant HTT protein (mHTT) and the preferential degeneration of the striatum. Previously in our lab, we identified Protein Kinase CK2 as an important kinase involved in the pathophysiology of HD. Specifically, the alpha prime catalytic subunit of CK2 (CK2α’) is upregulated in HD, and genetic depletion of CK2α’ in HD mice results in improved motor behavior, decreased mutant Htt aggregation, and improved neuronal function. Silmitasertib (CX-4945) is an FDA designated orphan drug that inhibits CK2. This study aims to investigate whether CX-4945 treatment ameliorates HD pathology. We treated prodromal and late symptomatic HD mice, and used a variety of immunohistochemical, biochemical, physiological and behavioral approaches. We found that CX-4945 presented benefits in the amelioration of HD pathophysiology in both treated groups. Importantly, we found CX-4945 decreased mHtt aggregation, increased DARPP-32 expression and excitatory synapse density, restored homeostatic astrocyte phenotypes and ameliorated neuroinflammation and microgliosis, altogether resulting in improved motor behavior. These results support CX-4945 as a strong candidate for a targeted therapy to treat HD.

## Introduction

Huntington’s disease (HD) is a neurodegenerative disease that presents with progressive motor, cognitive, and psychological impairments. HD is caused by a polyQ (CAG) repeat expansion in exon 1 of the huntingtin gene (*HTT*)(1). PolyQ expansion in the *HTT* gene results in misfolded mutant huntingtin protein (mHTT) and preferentially affects striatal GABAergic medium spiny neurons (MSNs) (1–3). Death of MSNs in the striatum leads to striatal tissue loss and progressive motor and cognitive impairments (3), although the mechanism behind the preferential loss of MSNs in HD is not well understood (4). Despite recent exciting developments in the implementation of strategies targeting HTT in the brain (5,6), there are currently no approved therapeutic treatments yet for HD.

Studies in both HD mouse models and humans have revealed that neuroinflammation and transcriptional dysregulation, along with mHTT aggregation, are key features in HD that precede MSN dysfunction and loss. Reactive gliosis is another major pathological hallmark of HD (7–10). MSN dysfunction in HD is often accompanied by the accumulation of hypertrophied microglia and astrocytes, increased astrocyte proliferation, altered astrocyte transcriptomic and proteomic profiles and dysregulation of glial homeostatic functions (11–13). It has long been recognized that MSN and glial cell alterations in HD contribute to the neuropathology and symptomatology of HD and are caused by both cell- and non-cell autonomous processes (14,15). However, the mechanisms that establish the reciprocal and functional interactions between MSNs and glial cells in HD are not fully understood (15–18).

Protein kinase CK2 is a serine/threonine kinase that plays essential roles in the regulation of neuroinflammation and protein aggregation (19–22). CK2 is composed of two beta regulatory subunits and two catalytic subunits CK2 alpha and CK2 alpha prime (CK2α and CK2α’) that differ in substrate specificity, expression abundance, and tissue distribution (19,20,23). CK2α is abundantly and ubiquitously expressed throughout the body while CK2α’ expression is more restricted to brain and testis (19,20,23). Previous work in our lab has shown that CK2α’ is preferentially upregulated in cell and mouse models of HD and in the striatum of HD patients (24). CK2α’ is specifically upregulated in MSNs, and its accumulation correlates with the onset of mHtt aggregation and motor deficits. Importantly, we showed that HD mice happloinsufficient for CK2α’ decreased mHtt aggregation, restored the transcriptional profile of both neurons and astrocytes, and improved MSNs synaptic function and motor behavior (25). Collectively, these genetic studies established CK2 as a promising therapeutic target and justified further investigation of pharmacological inhibitors of this kinase as potential treatment strategies.

A number of CK2 inhibitors have been developed, and notably, CX-4945 (Silmitasertib) and CIGB-300 have advanced to clinical evaluation in cancer patients (26–29), owing to the frequent upregulation of CK2 in several malignancies (23,30,31). CX-4945 has demonstrated a favorable safety profile in humans and has received FDA orphan drug designation. Although CX-4945 has been extensively used in preclinical cancer studies, its therapeutic efficacy in non-oncology contexts remains limited and often associated with in vitro manipulations. We and others have shown that CK2 is also induced in other neurodegenerative diseases such as spinal and bulbar muscular atrophy (SBMA), Alzheimer’s (AD) and Parkinson’s disease (PD) (23,24,32–35). Importantly, a recent study in a mouse model of AD reported that CX-4945 treatment reduced pathological features and ameliorated behavioral deficits (36). This recent study opens the possibility of repurposing CX-4945 as a therapeutic strategy for other neurodegenerative diseases, including HD.

In this study we explored the therapeutic effects of CX-4945 in the treatment of HD using the zQ175 mouse model (37). We provided CX-4945 by gavage administration at both prodromal and symptomatic ages and found the amelioration of several HD-associated features including decreased mHtt aggregation, improved striatal synaptic density and activity, decreased neuroinflammation, and improved astrocyte homeostatic functions, overall resulting in improved motor behavior. These results further support the use of CK2 inhibition as a promising therapeutic strategy for the treatment of HD.

## Materials and Methods

### Mouse lines

This study utilized a full-length humanized knock in exon 1 mouse model (zQ175) (37). This chimeric mHtt contains approximately 165 CAG repeats, recently analyzed in our mouse colony (25), and the human poly-proline region. This mouse model was chosen due to its slower and more realistic disease progression compared to more rapid transgenic models. Heterozygous zQ175 mice were obtained by crosses with WT (C57Bl/6J). C57Bl/6J mice were also used as controls. Mice were obtained from a total of 28 litters with an average of 7 pups. WT littermates were used when possible. Mice were treated at 5 months and 11 months respectively. Sample sizes are described throughout the study for each individual analysis. Cohorts were sex balanced when possible, however sex differences have not been reported in zQ175 mice or in CK2 depleted mice (25,37. All animal care and sacrifice procedures were approved by the University of Minnesota Institutional Animal Care and Use Committee (IACUC) in compliance with the National Institutes of Health guidelines for the care and use of laboratory animals under the approved animal protocol 2011-38628A.

### Immunofluorescence

Animals were anesthetized with Avertin (250 mg/kg Tribromoethanol) and perfused intracardially with tris-buffered saline (TBS) (25 mM Tris-base, 135 mM Nacl, 3 mM KCl, pH 7.6) supplemented with 7.5 mM heparin. Brains were dissected, fixed with 4% PFA in TBS at 4◦C for 4–5 days, cryoprotected with 30% sucrose in TBS for 4–5 days and embedded in a 2:1 mixture of 30% sucrose in TBS:OCT (Tissue-Tek), as previously described(24). Brains were cryo-sectioned into 16 µm-thick coronal sections, washed and permeabilized in TBS with 0.2% Triton X-100 (TBST). Three slices per animal at intervals of approximately 0.2 mm were chosen per animal. Slices were blocked for one hour in 5% normal goat serum in TBST, or bovine serum albumin for goat antibodies in PBST. Primary antibodies were diluted in 5% NGS in TBST or 5% BSA in PBST for goat antibodies. The primary antibodies that were utilized and accompanying dilutions are: GFAP (chicken, ab5541, 1:2000), S100B (rabbit, Abcam ab41548, 1:500), EM48 (mouse, mab5374, 1:500), GFP (chicken, Abcam AB13970, 1:1000), LAMP2 (rat, Thermo-Fisher, MA1-165, 1:100), Darpp32 (rat, R&D Systems MAB4230, 1:500), Iba1 (goat, Wako 011-27991, 1:500), cFos (Rabbit, Abcam ab190289, 1:500), MAP2 (Chicken, Abcam ab5392, 1:500) VGLUT1 (Guinea Pig, Millipore-Sigma AB5905, 1:500), VGLUT2 (Guinea Pig, Millipore-Sigma AB2251-I, 1:1000), PSD-95 (Rabbit, Thermofisher 51-6900, 1:500).

### Immunoblotting and cytokine profiler array

Animals were anesthetized with Avertin (250 mg/kg Tribromoethanol) and perfused intracardially with tris-buffered saline (TBS) (25 mM Tris-base, 135 mM Nacl, 3 mM KCl, pH 7.6) supplemented with 7.5 mM heparin. Microdissection under a dissecting microscope was performed to isolate the striatum. Striatum was homogenized in cell lysis buffer (25 mM Tris-HCl pH=7.4, 150 mM NaCl, 1% Triton, 1 mM EDTA, 0.1% SDS), for 60 seconds using a tissue grinder and pestle tip. 50 ug of protein was loaded on to BioRad 4-20% stain free gel and run for 2 hours at 80 V, or until loading buffer reached the bottom of gel. Gels were blocked with 5% non-fat powdered milk in TBST for 1 hour. Primary antibodies were diluted in 5% non-fat powdered milk in TBST overnight at 4 degrees C. Gels were imaged on an Amersham ImageQuant 800 Western Blot Imaging System. The primary antibodies that were utilized and accompanying dilutions are: EM48 (mouse, mab5374, 1:500), Darpp32 (rat, R&D Systems MAB4230, 1:500), GAPDH, (mouse, Santa-Cruz sc-365062, 1:5000). Mouse cytokine proteome profiler kit was purchased from Biotechne (ARY006) and used following manufacturer specifications using protein extracts obtained as described above.

### Imaging and image analysis

Image collection and analysis was performed as previously described in Brown et al. 2024. Images were collected using either an epifluorescent microscope (Echo Revolve) or a confocal microscope (Lecia Stellaris 8). On the Echo Revolve, 1.25x images have a 0.4467 pixel/µm resolution and 10x images have a 3.569 pixel/µm resolution. All Echo images have a 2732 × 1948 pixel size. Stellaris 10x images have a 0.17 pixel/µm resolution, 512 × 512 pixel size, and a z-stack encompassing the entire 16 µm was acquired. 63x images have an 856 x 856 pixel size, X pixel/µm resolution, and z-stack encompassing the entire 16 µm slice thickness was acquired. For 10x images, 3 images per slice were acquired with each containing three images corresponding to the dorsomedial, dorsolateral and centromedial striatum respectively. Images were analyzed in their entirety for all analyses. Image settings including laser intensity, laser gain and binning were set using control samples for each experiment. To ensure no overlapping occurred, images were taken at least 100 µm apart. Analyses were performed as previously described in Brown et al. 2023(7).

### Cell counting and puncta analysis

For cell counting, the call counter plugin from FIJI was used and cells were counted manually. For analyzing puncta, such as mHtt aggregates or colocalized puncta, the puncta analyzer plugin from FIJI was used. 10x Stellaris images with a 16 µm z-dimension were used. Counts from all 3 images per slice were averaged and animal averages were reported. Images were blinded using an automated blinding macro, and image analyses were done blind to treatment group and genotype. Analyses were performed as previously described in Brown et al. 2023(7).

### Colocalization analysis

10X Stellaris images for colocalization analysis and processed on the Imaris image editing software. Colocalization channels were created using consistent thresholds set by the positive control for each experiment. Colocalization channels were created, and FIJI puncta analyzer was used to count colocalized puncta.

### Calcium Imaging and Analysis

Calcium imaging on astrocytes was performed as previously described (16). HD and WT mice were bilaterally injected with .5uL of AAV5-GFAP-GcAMP6, previously used in (38,39), in the striatum. 2 weeks post injection, mice were rapidly decapitated under isoflurane and 350 µm brain slices were obtained using a vibrating blade microtome (Leica VT 1000S), in sucrose cutting solution. Slices were incubated in aCSF for 30 minutes at 37°C and for 60 minutes at room temperature. Imaging was performed using a Leica Stellaris 8 with a 25x water 2-photon objective. Recordings were performed for 15 minutes with 5 minutes of baseline recording. After 5 minutes baseline, either 50 mg/mL CX-4945 or saline solution were added. 4 animals per treatment group were utilized with a minimum of 30 cells per group. Specific sample sizes are reported in figure legends. Slices that contained less than 4 GcAMP expressing cells were excluded. For data analysis, AqUA, a freely available astrocyte calcium imaging analysis was used.

### Behavior

Animal behavior experiments were conducted in the University of Minnesota Mouse Behavior core. Males were always tested before females, and experimenter was blind to genotype and treatment group. For the *beam walk test*, animals were first acclimated during a 30-minute period on the room, then the beam walk task was conducted with three training days consisting of four trials per animal on the medium square beam. Animals were run in groups 5-7 to ensure an inter trial interval of 15 minutes or less. On test day, 2 trials for the beam were performed. Small round beam was 91cm long with a 10mm diameter. Trial time, number of foot slips, and number of stops were recorded.

For spatial recognition *Y-Maze*, after 30-minute acclimation, mice were placed in a Y-maze with one arm blocked off for 10 minutes. Mice were then returned to their home cages. After 1 hour, mice were placed back in the Y-maze with the covered arm open for 5 minutes. Time spent in each arm as well as total distance traveled was recorded. Animals were tested in groups of 5-7 to ensure a 1-hour rest interval.

*Open field* task was performed by placing mice in an open field chamber for 30 minutes, after a 30-minute acclimation period. Total distance traveled, time spent in center, and time spent in surround were recorded.

*Foot fault task* was performed by placing mice in a foot fault chamber after a 30-minute acclimation period. Number of foot faults and total distance traveled was recorded by the Foot Fault Scan 2 software.

Anymaze (AnyMaze, Wood Dale, IL) software was used for all behavior tasks except for foot fault. All behavioral assays and animal handling were approved by the University of Minnesota Institutional Animal Care and Use Committee (IACUC) in compliance with the National Institutes of Health guidelines for the care and use of laboratory animals under the approved animal protocol 2011-38628A.

### Experimental Design and Data Analysis

Data analysis and experimental design was performed as described above. Data from three brain slices per animal were averaged together to get an animal average, and data per animal was used to conduct statistical analyses. Sample sizes are included in each figure legend. Graphs represent Mean ± SEM, with statistical significance analyzed and displayed by Prism 9 software (GraphPad, San Diego, CA, USA). To test normality, the D’Agostino and Pearson test was used when sample size was greater than or equal to 8 and the Shapiro-Wilk test was used when sample size was less than 8. Normally distributed data were compared with t-test (two-tailed), ordinary one-way ANOVA with Tukey’s multiple comparisons test or two-way ANOVA with Sidak’s multiple comparison test were used when appropriate. Acceptable significance was p ≤ 0.05. n.s. indicates the comparison did not reach ≥ 0.05.

## Results

### CX-4945 treatment in HD mice decreased mHtt aggregation and spiny neuron dysfunction

We conducted CX-4945 treatments in both prodromal (5 months old) and symptomatic (11 months old) heterozygous zQ175 animals (referred hereafter as HD) and WT controls using gavage administration twice a day for 30 days with a 50 mg/kg dose (Figure 1A). The chosen dose was based on previous preclinical studies in cancer mouse models showing both safety and efficacy (40,41). We confirmed that the treatment was safe, as no significant loss in body weight was observed over the 30-day treatment period; importantly, none of the mice approached the established toxicity threshold of falling below 80% of their initial body weight (Figure 1B, C). We next assessed whether CX-4945 could ameliorate neuropathology in HD mice by evaluating two of the major pathological hallmarks of HD, mHtt aggregation and MSN dysfunction using both immunoblotting and immunofluorescence. In the prodromal group, we did not observe significant differences between saline and CX-4945 treated mice in mHtt aggregation (Figure 1D, E; G, I; K, L). However, in late symptomatic mice, CX-4945 treatment significantly decreased mHtt aggregation in neurons (Figure 1D, F; H, J; K, M). We previously demonstrated that CK2α’ is progressively upregulated in the striatum of HD mice and that its levels correlate positively with mHTT aggregation, with the strongest association starting at 12 months of age (25). Taken together, these findings suggest that CK2 inhibition might slow down the accumulation of aggregates over time. We next assessed the expression of DARPP-32, a marker of MSNs. DARPP-32 is one of the first MSNs markers to show downregulation and it is considered a sign of MSNs transcriptional dysregulation and dysfunction in HD (42,43). We found that CX-4945 did not affect the expression of DARPP-32 in the prodromal HD treated mice (Figure 2A, C, E, G) but significantly increased DARPP-32 levels in the symptomatic group (Figure 2B, D, F, H). This indicates that CK2 inhibition may exert stronger neuroprotective or restorative effects once MSN pathology has reached a more advanced state, aligning with the idea that late-stage molecular abnormalities may be more dependent on CK2-driven mechanisms.

**Figure 1.**
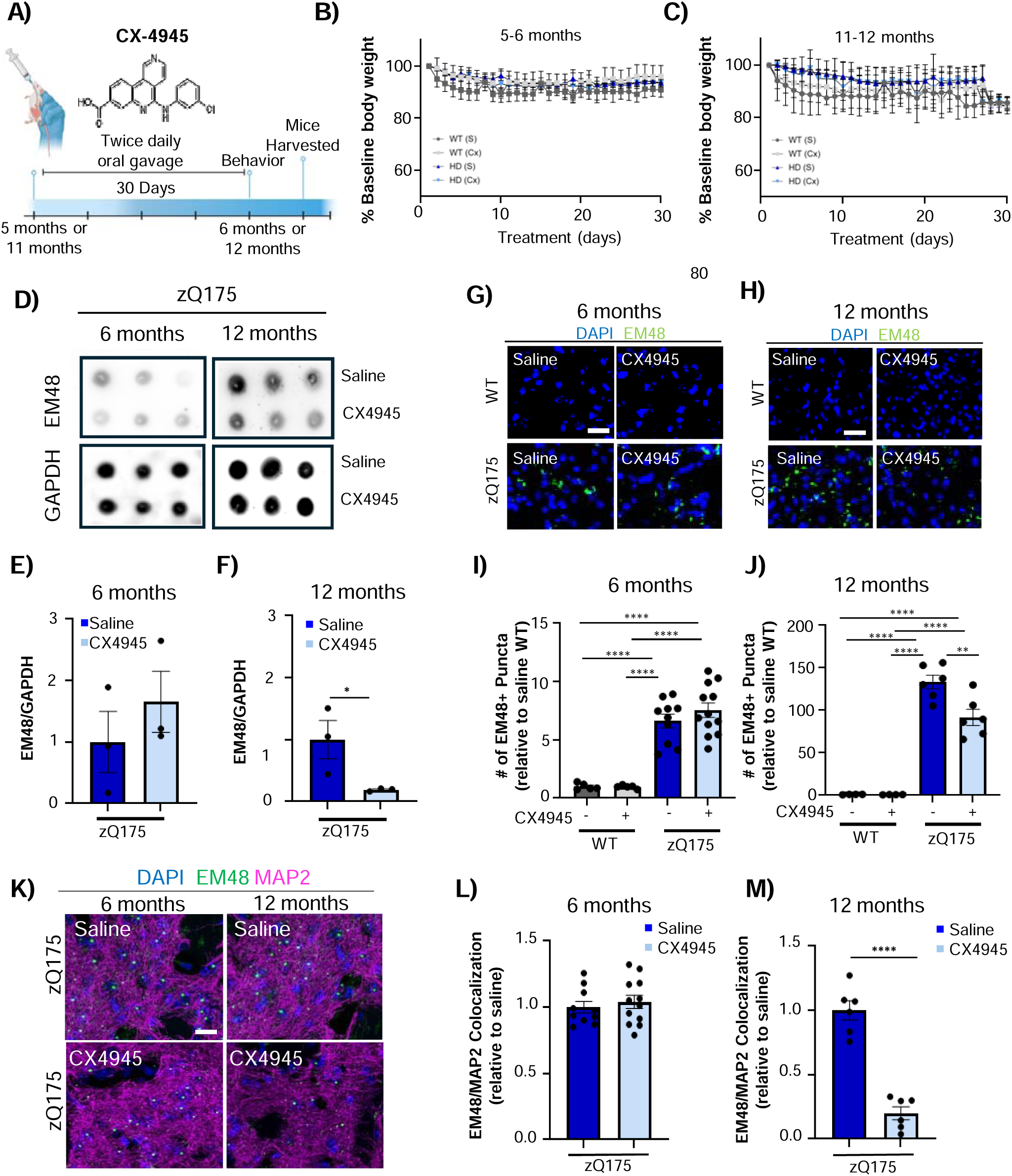
CX-4945 decreases mHTT aggregation in symptomatic zQ175 mice. A) Schematic representation of experimental design for CX4945 treatment. B, C) Average bodyweight measurements for prodromal (B) and symptomatic (C) mice during 30-day oral gavage treatment. D) EM48 and GAPDH dot blots from striatal protein extracts of 6-month and 12-month CX-4945 and saline treated zQ175 mice. E, F) EM48 protein levels relative to GAPDH from images in D (n=3 mice/group). G, H) EM48 immunostaining as a marker of mHtt aggregates and DAPI (stains nuclei) in dorsal striatum sections of 6-month (G) and 12-month (H) WT and zQ175 mice treated with CX4945 or saline. Scale bar, 50 μm. I, J) Quantification of EM48+ puncta per 0.4 mm^2^ in the striatum of 6-month (I) and 12-month (J) WT and zQ175 mice treated with CX4945 or saline (n=5-12 mice per group) analyzed from images in G and H respectively. Data was normalized to number of DAPI+ cells and relativized to saline controls. K) EM48 and MAP2 immunostaining in dorsal striatum sections of 6-month and 12-month mice treated with CX4945 or saline neurons with DAPI (stains nuclei) in striatum of 6-month and 12-month CX4945 or saline. Scale bar, 5 μm. L, M) Quantification of EM48+ puncta colocalized with MAP2+ neurons in 6-month (L) and 12-month (M) zQ175 mice treated with CX4945 or saline (n=10-12 mice/group for L, n=6mice/group for M) from images in K. Data was normalized to number of DAPI+ cells and relativized to saline controls. Error bars denote mean ± SEM. Student’s unpaired t-test [E) t=.9292 dF=4, F) t-3.204 dF=4, L) t = .5951 dF = 20, M) t = 8.800 dF = 10] and ordinary one-way ANOVA with Tukey’s multiple corrections [I) F = 29.99 dF = 3, J) F = 69.96 dF = 3]. \**p* < 0.05, \*\**p* < 0.01, *p*<0.001, \*\*\*\**p* < 0.0001.

**Figure 2.**
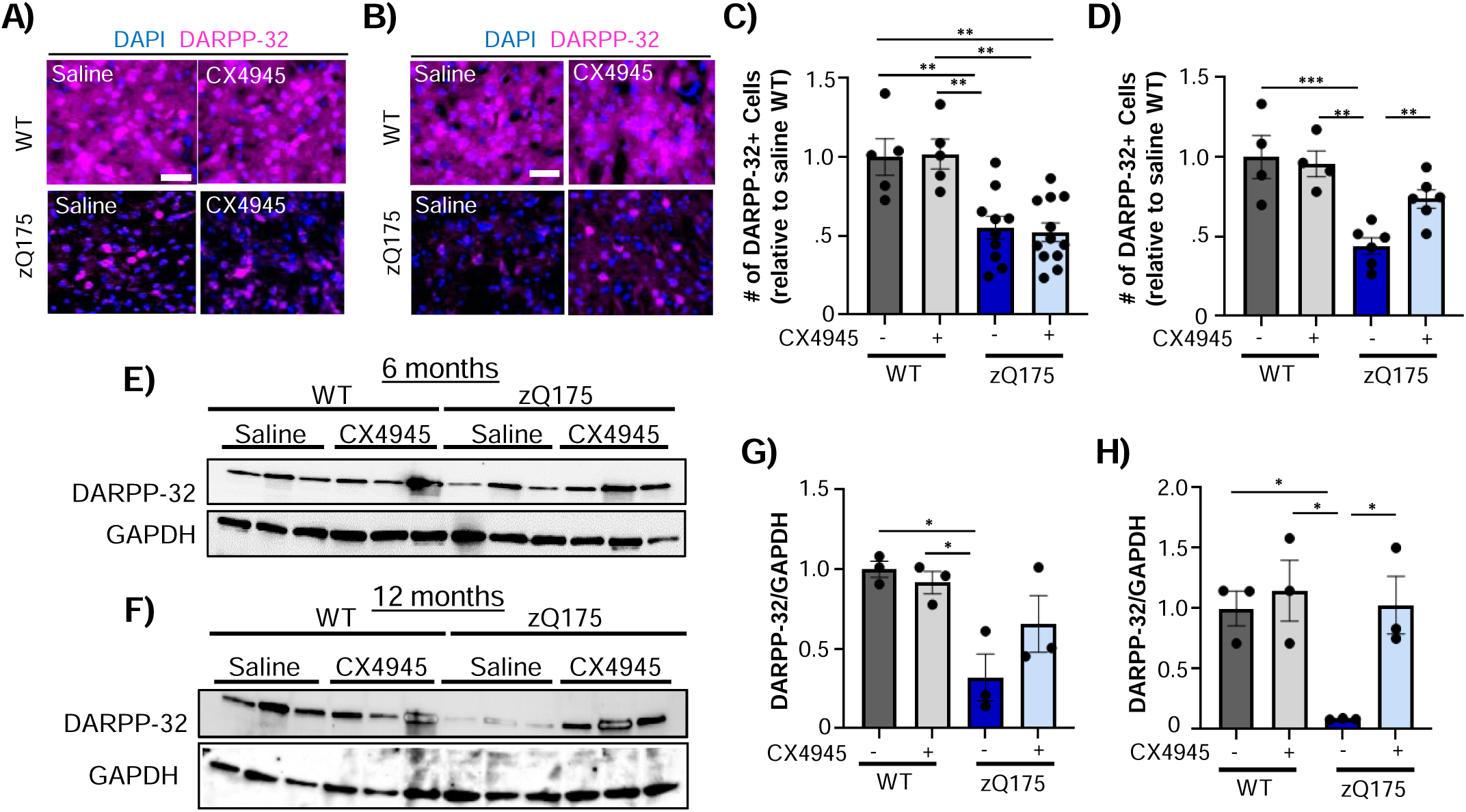
CX-4945 increased the number of DARPP-32+ cells in zQ175 mice. A, B) Immunostaining of DARPP-32+ MSNs in the dorsal striatum of 6-month (A) and 12-month (B) WT and zQ175 mice treated with CX4945 or saline. DAPI stains nuclei. Scale bar, 50 μm. C, D) Quantification of number of DARPP-32+ cells per 0.4 mm^2^ in the dorsal striatum of 6-month (A) and 12-month (B) WT and zQ175 mice treated with CX4945 or saline (n=4-12 mice/group) analyzed from images in A and B respectively. Data was normalized to number of DAPI+ cells and relativized to saline WT. E, F) DARPP-32 immunoblotting from striatum protein extracts of 6-month (E) and 12-month (F) mice treated with CX4945 or saline. GAPDH was used as a loading control. G, H) DARPP-32 protein levels relative to GAPDH analyzed from images in E and F respectively (n=3mice/group). Error bars denote mean ± SEM. Ordinary one-way ANOVA with Tukey’s multiple corrections [C) F = 10.48 dF = 3, D) F = 11.21 dF = 3, G) F = 6.164 dF = 3, H) F=6.857 dF=3]. \*\**p* < 0.01, *p*<0.001, \*\*\*\**p* < 0.0001.

To better assess whether CX-4945-mediated changes in DARPP-32 translated into an overall improvement in MSNs’ health we analyzed striatal synapse density and neuronal activity. Loss of cortico-striatal excitatory synapses has been widely reported in various HD mouse models, especially at older ages (44–46). Loss of synapses is accompanied by the depletion of c-fos expression, an immediate early gene induced by neuronal activity (47,48), indicative of the associated loss of neuronal function. We therefore investigated the effect of CX-4945 on cortico-striatal synapse density by assessing the colocalization of the pre-synaptic marker PSD-95 and the cortical-derived vesicular transporter VGLUT1 using confocal imaging. As previously reported, no significant changes were observed in the levels of cortico-striatal synapses at 6 months old across genotypes, and no changes were observed with CX-4945 treatment (Figure 3A-D). For the symptomatic groups, we observed a significant depletion of cortico-striatal synapses in saline treated HD mice compared to WT that was driven by the depletion of both PSD-95 and VGLUT1 expression (Figure 3E-H). Importantly, CX-4945 significantly increased synapse density of cortico-striatal synapses in symptomatic treated mice (Figure 3 E-H). To determine whether an increase in synapse density was translated into enhanced neuronal activity we utilized immunofluorescent detection of cFos. We observed a modest reduction in c-fos expression in the saline treated HD-mice compared to WT at 6 months of age, which became more pronounced by 12 months (Figure 3I-J). Treatment with CX-4945 significantly increased the number of cFos+ cells relative to saline-treated HD mice in both the prodromal and late symptomatic cohorts. Notably, the rescue effect at 12 months was substantially greater than that observed at 6 months, indicating an enhanced responsiveness to CK2 inhibition at later disease stages (Figure 3I-J).

**Figure 3.**
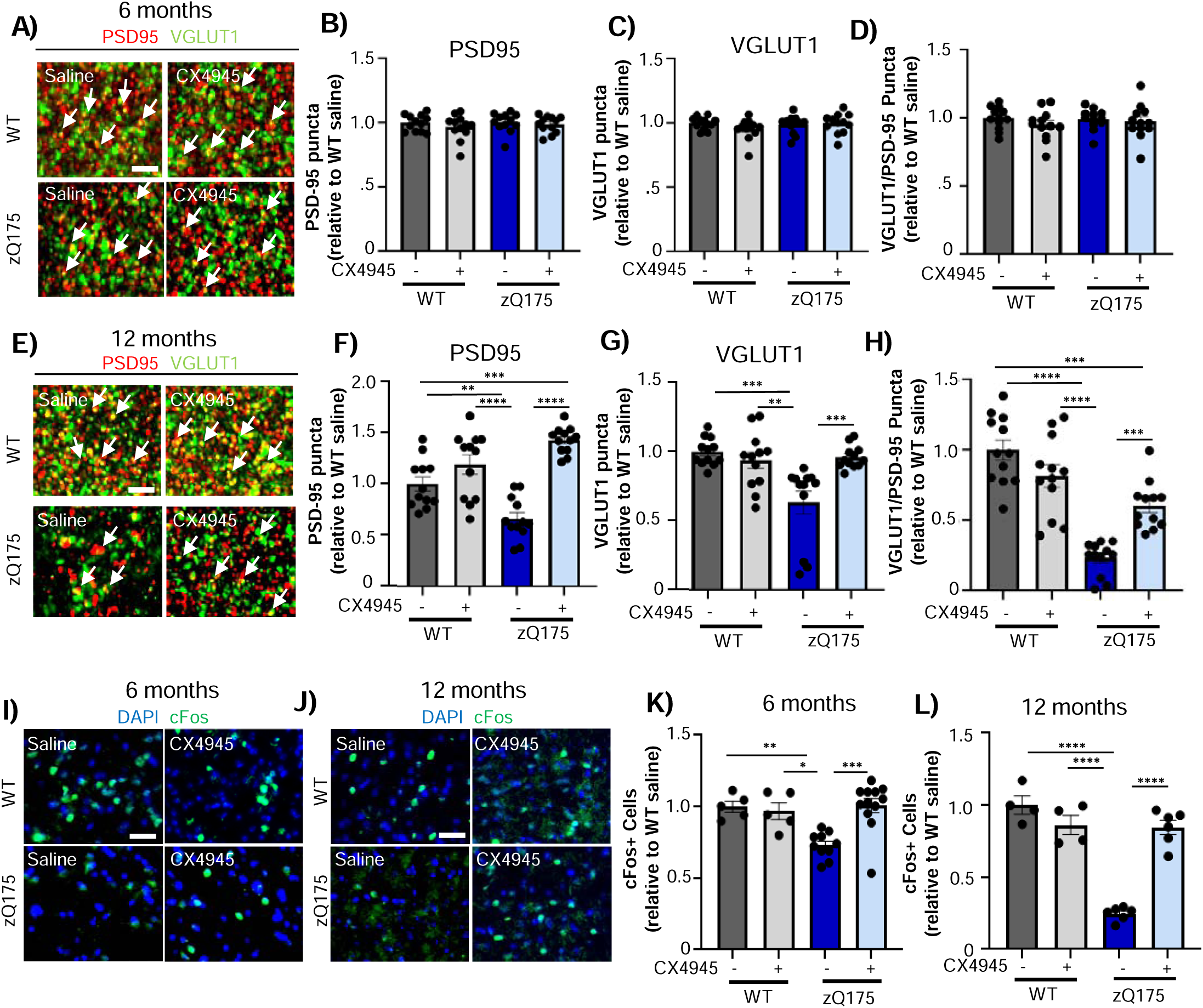
CX-4945 treatment increases cortico-striatal synaptic density and neuronal activity in zQ175 mice. A) PSD-95 and VGLUT1 immunostaining in the dorsal striatum of 6-month WT and zQ175 mice treated with CX-4945 or saline. Highly contrasted images are shown for representation of colocalized puncta indicated by white arrows. Scale bar, 2 μm. B-D) Quantification of PSD-95 puncta (B), VGLUT1 puncta (C), and colocalized PSD-95/VGLUT1 puncta (D) from images in A (n=12 images/group corresponding to 3 slices/mouse and 4 mice/group. Slice averages/mouse are shown). E) PSD-95 and VGLUT1 immunostaining in the dorsal striatum of 12-month WT and zQ175 mice treated with CX-4945 or saline. Highly contrasted images are shown for representation of colocalized puncta indicated by white arrows. Scale bar, 2 μm. F-H) Quantification of PSD-95 puncta (F), VGLUT1 puncta (G), and colocalized PSD-95/VGLUT1 puncta (H) from images in A (n=12 images/group corresponding to 3 slices/mouse and 4 mice/group. Slice averages/mouse are shown). I, J) cFos immunostaining in the dorsal striatum of 6-month (I) and 12-month (J) mice treated with CX4945 or saline. Scale bar, 50 μm. K, L) Quantification of cFos+ cells per 0.4 mm^2^ in the dorsal striatum of 6-month (K) and 12-month (L) mice treated with CX4945 or saline (n=4-12 mice/ group) analyzed from images in I and J respectively. Data was normalized to number of DAPI+ cells and relativized to saline WT. Error bars denote mean ± SEM. Ordinary one-way ANOVA with Tukey’s multiple comparisons [B) F = .1829 dF = 3, C) F = .2514 dF = 3, D) F = .5828 dF = 3, F) F = 22.85 dF = 3, G) F = 9.597, dF = 3, H) F = 28.96 dF = 3. \**p* < 0.05, K) F = 9.549 dF = 3, L) F = 52.79 dF = 3]. \**p* < 0.05, \*\**p* < 0.01, *p*<0.001, \*\*\*\**p* < 0.0001.

### CX-4945 decreased striatal inflammatory cytokine production and microglial phenotypes in HD treated mice

CK2 has been previously associated with the regulation of inflammation in different contexts, including neurodegenerative diseases (22,49,50). Increased expression of inflammatory cytokines and abnormal microglial phenotypes in the striatum are additional pathological hallmarks of HD (51–54). These microglial alterations are considered a secondary response to neuronal dysfunction. Thus, we hypothesized that the benefits in neuronal activity and synapse density mediated by CX-4945 treatment in HD mice may also influence microglial phenotypes and neuroinflammation. We therefore investigated the impact of CX-4945 on reactivemicrogliosis and cytokines production. Using Iba1, a marker for activated microglia, we assessed the number of Iba1+ cells in prodromal and late symptomatic CX-4945 treated HD mice. We observed an increase in Iba1+ cells in both HD groups compared to WT mice, consistent with microgliosis, however CX-4945 did not significantly affected Iba1+ cells in either age group (Figure 4A-D). While reactive microglia proliferate in HD and other neurodegenerative diseases, they also undergo morphological changes which are more strongly related to the reactive state of microglia. Homeostatic microglia or surveilling microglia have more branches and branch points while amoeboid microglia have fewer branches and are associated with a more reactive phenotype and the production of inflammatory cytokines (51,55). We assessed microglia morphology using high magnification confocal microscopy in both prodromal and symptomatic groups (Figure 4E-H). We only found a significant reduction in the number of branches in saline-treated HD mice at symptomatic age, consistent with an increased ameboid phenotype (Figure 4F, H). Importantly, CX-4945 treatment decreased the microglial ameboid phenotype by increasing the number of microglial branches compared to saline-treated HD mice (Figure 4F, H).

**Figure 4.**
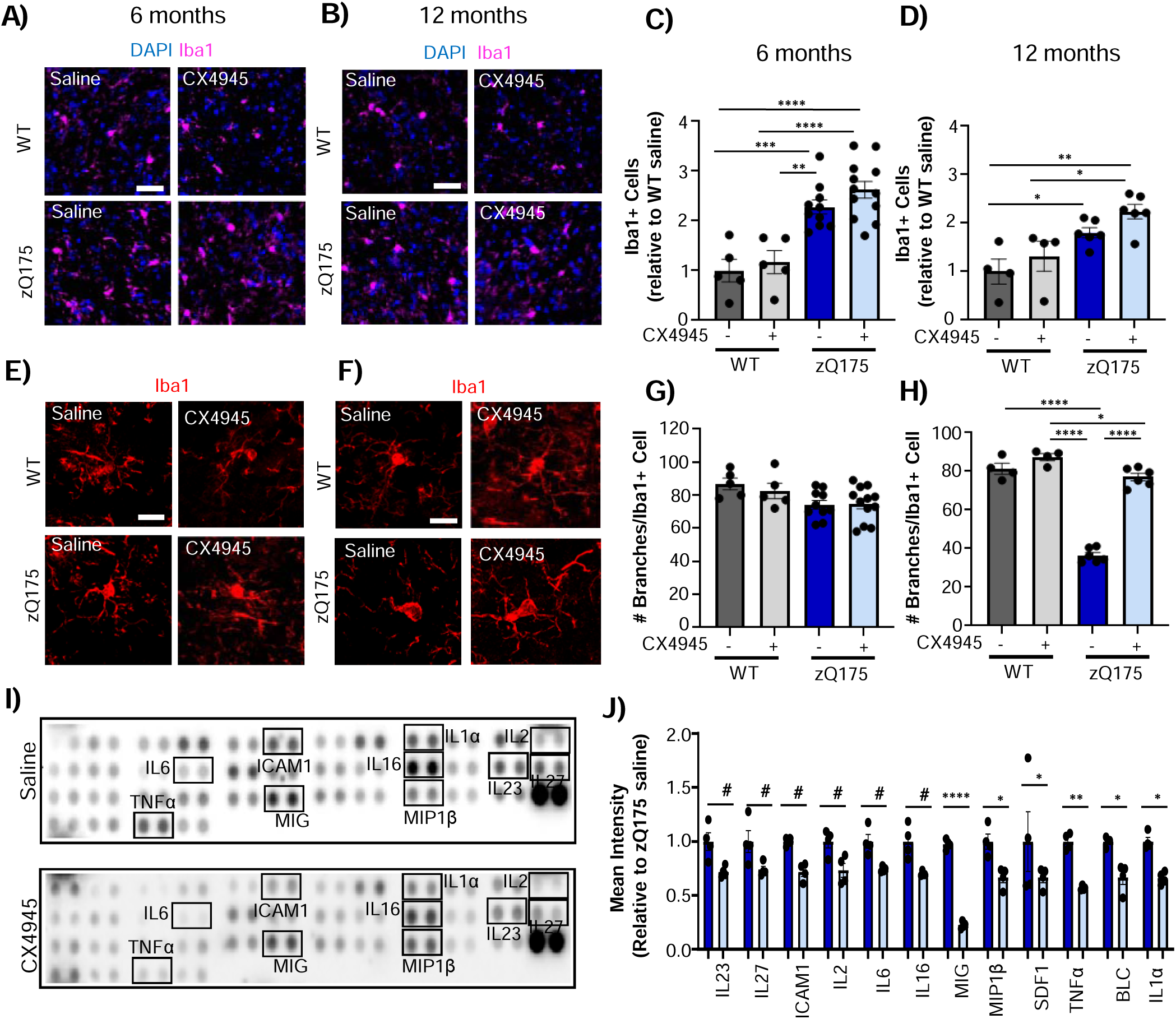
CX-4945 treatment restores microglia morphological abnormalities and reduces inflammatory cytokines in zQ175 mice. A, B) Iba1 immunostaining in the dorsal striatum of 6-month (A) and 12-month (B) WT and zQ175 mice treated with CX4945 or saline. Scale bar, 50 μm. C, D) Quantification of Iba1+ cells per 0.4 mm^2^ in the dorsal striatum of 6-month (C) and 12-month (D) mice treated with CX4945 or saline (n=4-12 mice/group) analyzed from images in A and B respectively. Data was normalized to number of DAPI+ cells and relativized to saline WT. E, F) Representative images of individual Iba1+ cells in the dorsal striatum of 6-month (E) and 12-month (F) mice treated with CX4945 or saline. Scale bar, 5 μm. G, H) Quantification of the number of branches per Iba1+ microglia in 6-month (G) and 12-month (H) WT and zQ175 mice treated with CX-4945 or saline from images in E and F respectively (n=4-12 mice/group). I) Representative mouse cytokine array panel from striatum protein extracts from 12-month zQ175 mice treated with CX-4945 or saline. Cytokines showing significant or trending changes between CX-4945 and saline treated mice are highlighted in black. J) Relative cytokine levels normalized to internal loading control and analyzed from images in I (n=4 mice/group). Error bars denote mean ± SEM. Ordinary one-way ANOVA with Tukey’s multiple comparisons [C) F=16.78 F=3, D) F=7.717 dF=3, G) F=2.799 dF=3 H) F=146.2 dF=3], and two way ANOVA with Sidak’s post-hoc test [J) Interaction = 1.636 (dF = 11), Genotype = 129.0 dF = 1), Cytokine = 1.979 (dF = 11). #p<0.1, \**p* < 0.05, \*\**p* < 0.01, *p*<0.001, \*\*\*\**p* < 0.0001.

To determine whether changes in microglial phenotypes may have a functional impact in the production of inflammatory cytokines, we examined the profile of several pro- and anti-inflammatory cytokines in saline and CX-4945 treated HD mice. Utilizing a cytokine proteome profiling array, we measured the relative level of 40 different cytokines (Figure 4I, J, Supplementary Table 1). CX-4945 significantly reduced the relative level of 6 different pro-inflammatory cytokines (MIG, MIP1β, SDF1, TNFα, BLC and IL1α), with another 6 cytokines (IL23, IL27, ICAM1, IL2, IL6, IL16) trending towards a reduction in the CX-4945 treated mice (Figure 5 I, J). No cytokines either significantly increased or showed a trend towards increasing in the CX-4945 treated mice (Supplementary Table 1). The overall reduction in cytokine level in CX-4945 treated mice along with the amelioration of ameboid microglial phenotypes demonstrated an overall decrease in the striatal inflammatory state.

**Figure 5.**
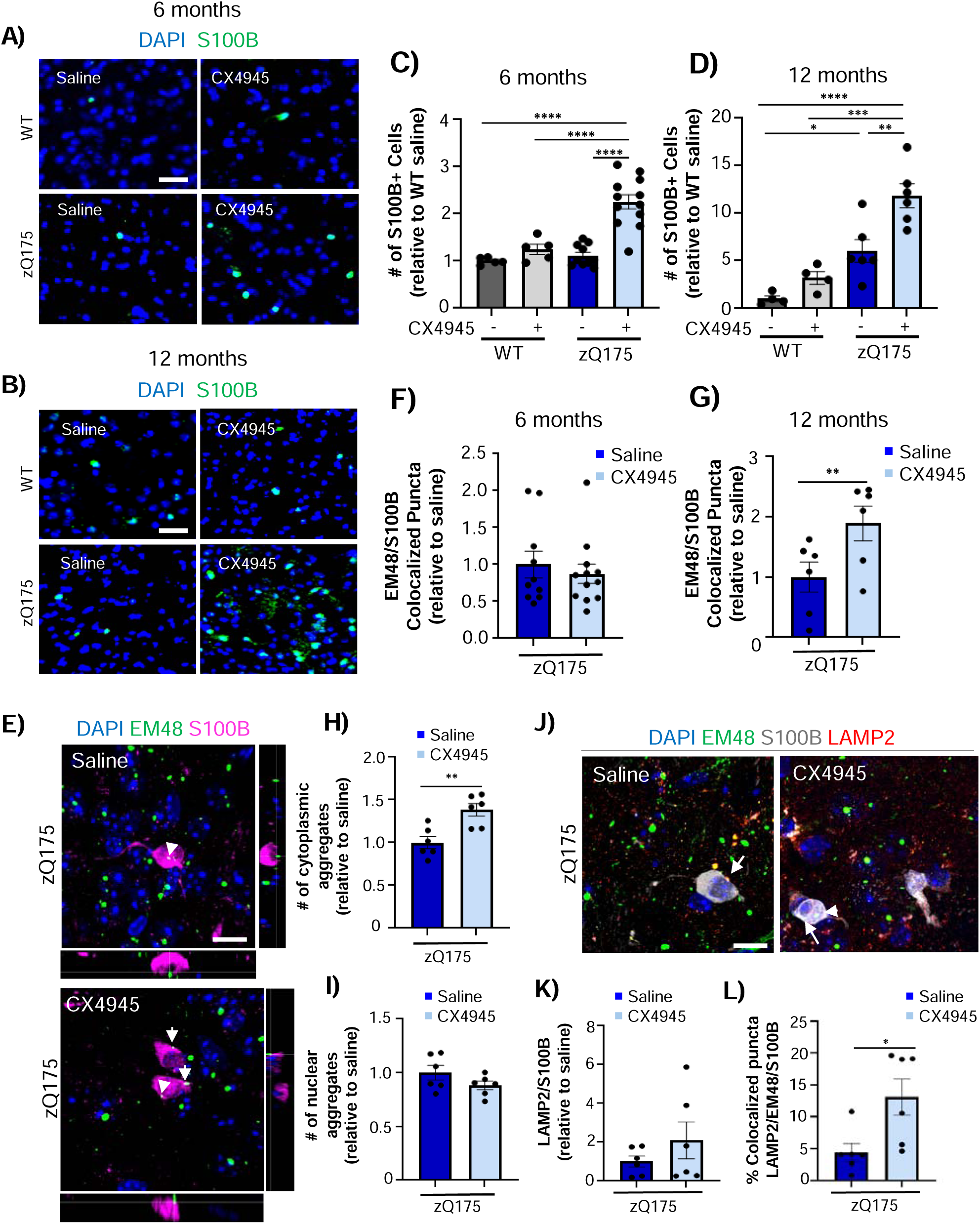
CX-4945 treatment increases S100B+ and restores phagocytic activity in zQ175 mice. A, B) S100B immunostaining in the dorsal striatum of 6-month (A) and 12-months (B) WT and zQ175 mice treated with CX4945 or saline treated. Scale bar, 50 μm. C, D) Quantification of S100B+ cells in the dorsal whole striatum of 6-month (C) and 12-months (D) Wt and zQ175 mice treated with CX4945 or saline (n=4-12 mice/group) analyzed from images in A and B respectively. Data was normalized to number of DAPI+ cells and relativized to saline WT. E) EM48+ and S100B+ immunostaining astrocytes within the dorsal striatum of 12-month zQ175 mice treated with CX4945 or saline. DAPI (stains nuclei) Scale bar, 5 μm. F, G) Quantification of EM48/S100B colocalized puncta in 6-month (F) and 12-months (G) zQ175 mice treated with CX4945 or saline (n=6-12mice/group). H, I) Quantification of cytoplasmic (H) and nuclear (I) EM48+ aggregates in S100B+ astrocytes in 12-month zQ175 mice treated with CX4945 or saline (n=6 mice/group) analyzed from images in E. J) EM48, S100B and LAMP2 immunostaining in 12-month zQ175 mice treated with CX4945 or saline. K) Quantification of LAMP2+ puncta in S100B+ astrocytes in 12-month zQ175 mice treated with CX4945 or saline (n=6 mice/group). Analyzed from images in J. L) Percent of colocalized EM48/LAMP2/S100B puncta analyzed from images in J (n=6 mice/group). Error bars denote mean ± SEM. Ordinary one-way ANOVA with Tukey’s multiple corrections [C) F= 24.49 dF=3, D) F=18.43 dF=3] and student’s unpaired t-test [F) t=0.5742 dF=17.24, G) t=2.341 dF=10, H) t=3.761 dF=10, I) t-1.556 dF=10, K) t=1.108 dF=5.918, L) t=3.121 dF=10]. \*\**p* < 0.01, *p*<.001, \*\*\*\**p* < 0.0001.

### CX-4945 treatment improved phagocytic and homeostatic astrocytic functions in HD mice

Given the close interplay between microglial activation, cytokines production, and astrocytic stress responses, we next investigated whether the modulation of inflammatory tone by CX-4945 impacted astrocytic reactivity. We performed immunofluorescence analyses in the dorsal striatum with S100B, an astrocytic reactivity marker (7,56). Contrary to what was expected we found a significant increase in the number of S100B+ astrocytes in CX-4945-treated HD mice in both prodromal and symptomatic groups (Figure 5A-D). This finding suggested an elevation in astrocytic reactivity. However, given the overall neuroprotective effects of CX-4945 observed in striatal pathology and neuroinflammation, this increase in S100B⁺ astrocytes were unlikely to reflect a detrimental response. Instead, we hypothesized that this response may indicate an adaptive or neuroprotective form of astrocyte activation associated with improved neuronal homeostasis. There’s growing evidence that S100B upregulation can be associated with increased astrocyte phagocytic activity (57,58) and therefore accumulation of S100B^+^ reactive astrocytes can exert neuroprotective effects in different contexts (59).

We next assessed whether mHtt aggregates differentially accumulate in S100B^+^ astrocytes and if such accumulation was impacted by CX-4945. We analyzed mHtt aggregate load in S100B^+^ astrocytes in prodromal and late symptomatic CX-4945 treated mice. In prodromal treated mice we did not observe a difference in the presence of mHtt aggregates in S100B+ astrocytes between saline and CX-4945 (Figure 5F). However, we observed an increase in the number of mHtt aggregates in S100B^+^ astrocytes in symptomatic mice treated with CX-4945 (Figure 5E, G). Interestingly, we observed that the increase in mHtt within S100B^+^ astrocytes was due to an increase in cytoplasmic aggregates, but not nuclear aggregates (Figure 5H, I). We hypothesized that this phenomenon could be mediated by an increase in astrocyte phagocytic activity, which takes place in cytoplasm. Similar to microglia, astrocytes are also known to engulf synapses and other debris in neurodegenerative diseases through LAMP2^+^ vesicles (60–62). To assess if astrocytes in CX-4945 present enhanced phagocytic activity we investigated the association between LAMP2 and mHtt aggregates in S100B^+^ astrocytes in symptomatic animals. While we did not observe a significant difference in the amount of LAMP2 within S100B^+^ astrocytes in either group (Figure 5J, K), we found a significant increase in the colocalization between LAMP2 and EM48 in S100B^+^ astrocytes of CX-4945 treated HD (Figure 5J, L). One possible explanation for these results is that CX-4945 facilitates the phagocytosis of Htt aggregates.

Notably, there is evidence for reciprocal regulatory feedback between S100B and CK2 where CK2 phosphorylates S100B and modulates its activity, while S100B upregulation activates CK2-dependent signaling cascades (59,63). CK2 and S100B also participate in Ca^²⁺^-dependent signaling and the regulation of stress responses in astrocytes and neurons (64–67). Previously, we showed that genetic deletion of CK2α’ in HD mice ameliorated transcriptional dysregulation of astrocytic genes related to glutamate uptake, phagocytosis, and Ca^²⁺^ signaling (25). Since phagocytosis and Ca^2+^ signaling are considered crucial astrocytic homeostatic functions, they are decreased in HD, and associated with CK2, we investigated whether CX-4945 also restored astrocytic Ca^²⁺^ signaling.

We injected prodromal and late symptomatic WT and HD mice with a Ca^²⁺^ sensor (GcAMP6f) under the control of GFAP promoter. Two weeks post-injection, brain slices were obtained, and viral expression specificity was confirmed, slices were then incubated with saline or CX-4945, and astrocytic Ca^²⁺^ transients were recorded *ex vivo* using 2-photon imaging (Figure 6A-C). We focused on assessing spontaneous astrocytic Ca^²⁺^ transients since previous studies have shown that spontaneous Ca^²⁺^ transients are significantly reduced in HD mice (16). First, we confirmed a reduction in the amplitude (dF/F0) and in the number of transients/min in saline-treated HD mice compared to WT in both prodromal and late symptomatic cohorts (Figure 6D-O). Both amplitude and transients/min were significantly increased in CX-4945-treated HD slices (Figure 6D-O). Taken together, these results suggest that enhanced expression of S100B^+^ astrocytes mediated by CX-4945 is associated with an improvement in astrocytic homeostatic functions of these astrocytes, especially in late symptomatic mice.

**Figure 6.**
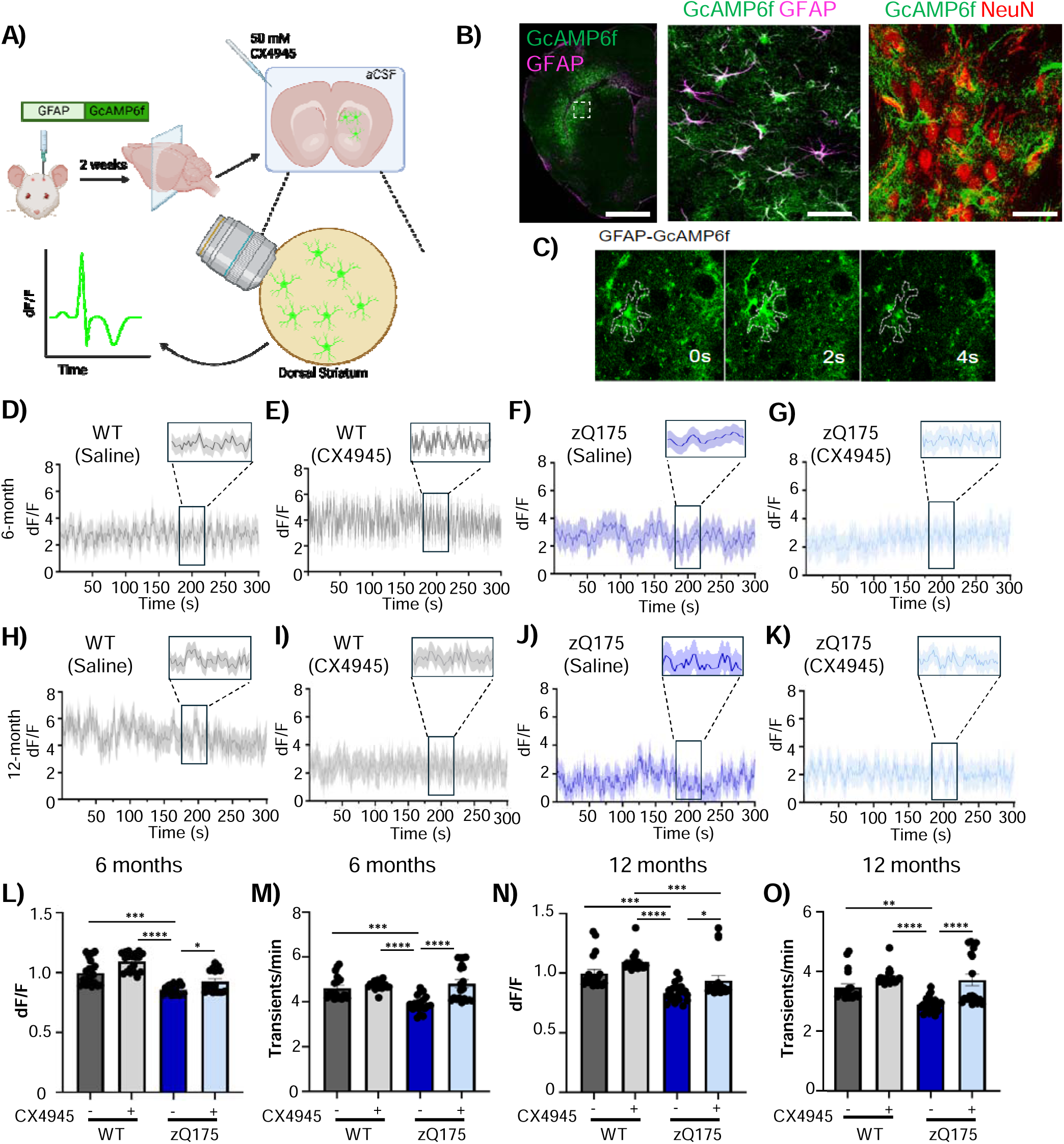
CX-4945 rescues Ca^2+^ signaling in astrocytes in zQ175 mice. A) Schematic of experimental design for the assessment of astrocytic Ca^2+^ signaling in the presence of CX-4945. B) Left panel corresponds to a brain coronal section of the dorsal striatum of WT mice injected with GFAP-GcAMP. Middle panel corresponds to GcAMP/GFP and GFAP immunostaining showing colocalized signal. Right panel shows NeuN and GcAMP/GFP immunostaining showing no colocalization. C) Representative image and of single astrocyte calcium spike between 0 and 4s. D-K) Representative GcAMP traces from individual astrocytes from the dorsal striatum in 6-month (D-G) and 12-month (H-K) WT and zQ175 brain slices incubated with CX4945 or saline. L, M) Amplitude (L) and frequency (transients/min) of Ca^2+^ events (M) in 6-month WT and zQ175 brain slices incubated with CX4945 or saline. Data was normalized to background fluorescence and relativized to saline treated WT mice (n=24-36 cells/group from a total of n=3 mice/group). N) Amplitude (N) and frequency (transients/min) of Ca^2+^ events (O) in 12-month WT and zQ175 brain slices incubated with CX4945 or saline. Data was normalized to background fluorescence and relativized to saline treated WT mice (n=24-36 cells/group from a total of n=3 mice/group). Error bars denote mean ± SEM. Ordinary one-way ANOVA with Tukey’s multiple comparisons [K) F = 29.98 dF = 3, L) F = 12.70 dF = 3, M) F = 15.82 dF = 3, N) F = 11.69 dF = 3]. \**p* < 0.05, \*\**p* < 0.01, p<0.001, \*\*\*\**p* < 0.0001.

### CX-4945 ameliorates motor deficits in prodromal and late symptomatic HD mice

We next investigated whether the effects mediated by CX-4945 in mHtt aggregation, synaptic density and function, inflammation, microgliosis, and homeostatic astrocyte functions translated into changes in HD related behaviors. We previously showed that genetic deletion of CK2α’ significantly rescued motor deficits in HD mice but did not rescue cognitive functions (25). Therefore, we focused our analyses on motor behaviors such as beam walk, foot fault test, and open field. In both prodromal and late symptomatic CX-4945 treated mice, we observed a significant decrease in the time mice needed to cross the beam and number of foot slips per quadrant in the beam walk test (Figure 7A-D). In the automated foot fault test, we observed notable variability in the distribution of performance that resulted in no significant differences in the overall mean number of foot faults between groups (Figure 7E). We then categorized animals into three bins corresponding to low (0–10), medium (11–30), and high (>30) numbers of foot faults per trial. With this analysis we found that saline-treated HD mice presented a higher percentage of high foot faults compared to CX-4945-treated HD mice (Figure 7F), suggesting that CX-4945 improved performance in the foot fault test. We also assessed motor performance in the open field test, but no significant differences were observed between groups or treatment in the prodromal or symptomatic groups (Figure 8G, H). Overall, our results showed that CX-4945 ameliorated motor deficits in HD mice.

**Figure 7.**
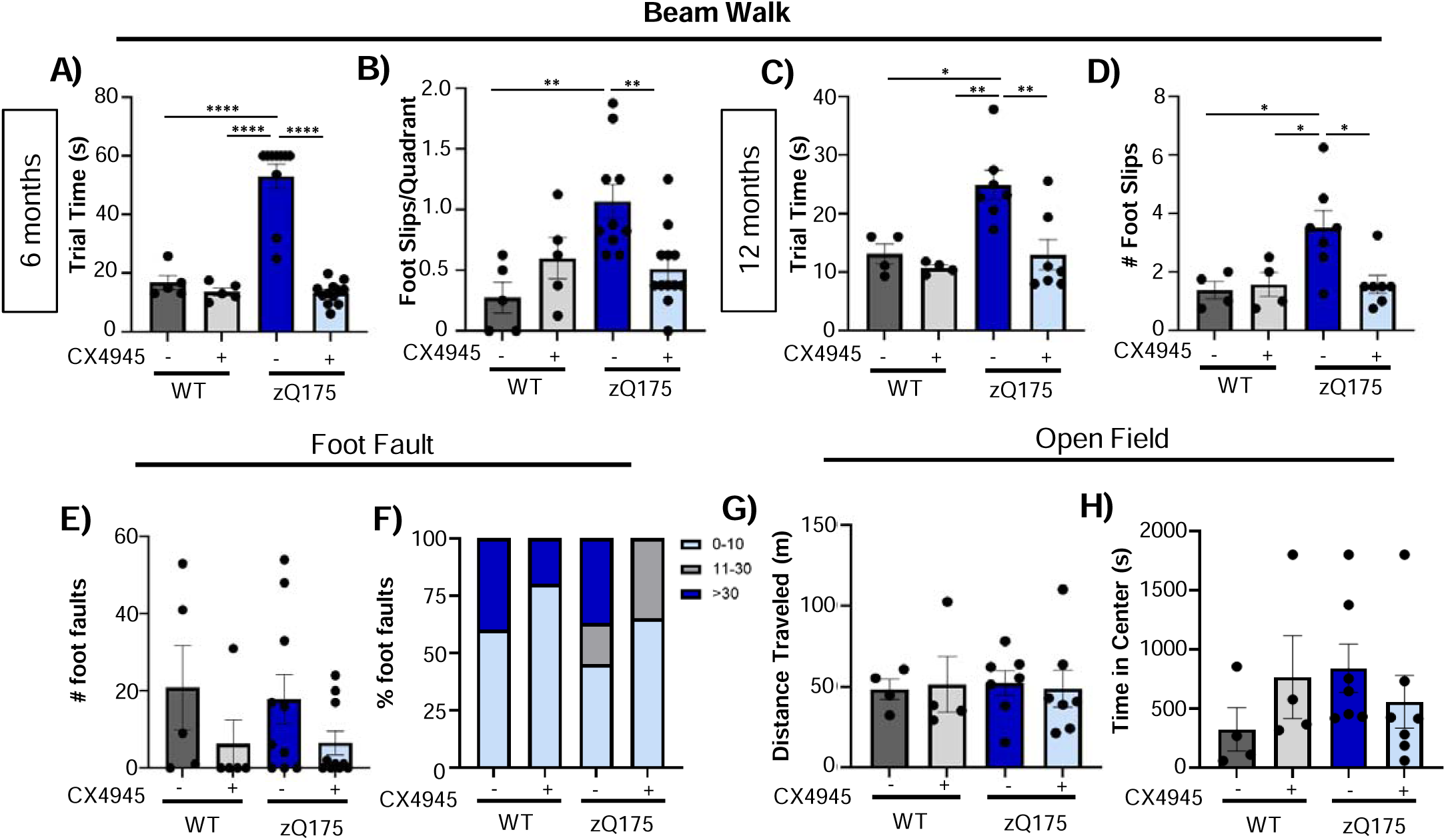
CX4945 ameliorates motor deficits in zQ175 mice. A) Average trial time to cross the beam in 6-month WT and zQ175 mice treated with CX4945 or saline (n=5-12 mice/group). B) Number of foot slips during beam crossing in 6-month WT and zQ175 mice treated with CX4945 or saline (n=5-12 mice/group). Number of foot slips is averaged per beam quadrant. If animal did not complete the beam after 60 seconds, the quadrant of the beam they ended the task in was recorded and foot slips were divided by number of quadrants completed. C) Average trial time to cross the beam in 12-month WT and zQ175 mice treated with CX4945 or saline (n=4-7 mice/group). D) Number of foot slips during beam crossing in 12-month WT and zQ175 mice treated with CX4945 or saline (n=4-7 mice/group). E) Average number of foot faults in 6-month WT and zQ175 mice treated with CX4945 or saline in the automated foot fault task (n=5-12 mice/group). F) Percentage distribution of animals presenting low (0–10), medium (11–30) or high (>30) foot faults per trial in 6-month WT and zQ175 mice treated with CX-4945 or saline (n=5-12 mice/group). G) Average distance traveled in open field task in 12-month WT and zQ175 mice treated with CX-4945 or saline (n=4-7 mice/group). H) Time spent in the center of the open field for 12-month zQ175 and WT mice treated with CX-4945 or saline (n=4-7 mice/group). Error bars denote mean ± SEM. Ordinary one-way ANOVA with Tukey’s multiple comparisons [A) F = 51.96 dF = 3, B) F = 6.433 dF = 3, C) F = 7.868 dF = 3, D) F = 5.195 dF = 3, E) F = 1.321 dF = 3 G) F = .02829 dF = 3 H) F=.1215 dF=3]. \**p* < 0.05, \*\**p* < 0.01, *p* < 0.0001.

## Discussion

CK2 has been previously identified as a potential therapeutic target in several neurodegenerative diseases including HD (68–70). The therapeutic potential in HD is supported by previous studies showing that genetic depletion of one of the catalytic subunits of CK2 (CK2α’) in zQ175 mice decreased mHtt aggregation, improved synaptic density and neuronal function, and ameliorated neuroinflammation and motor deficits. Unfortunately, earlier attempts to pharmacologically inhibit CK2 in HD cell models yielded conflicting results in ameliorating HD-like phenotypes, largely due to the poor selectivity and high toxicity of first-generation CK2 inhibitors such as TBB or Emodin (71,72). These limitations had precluded the investigation of the efficacy of these CK2 inhibitors in HD mouse models. However, the development of more selective and less toxic inhibitors, such as CX-4945, recently designated an FDA orphan drug, its oral bioavailability and brain blood barrier penetration, and its successful use in animal models of AD (36) have made this compound a more attractive candidate for HD therapy.

In our study we used the zQ175 knock-in HD mouse model and tested the efficacy of CX-4945 in the amelioration of striatal pathology and motor deficits at both prodromal and late symptomatic ages. We found that CX-4945 decreased mHtt aggregation in the symptomatic group but not in the prodromal group where Htt aggregation just started to accumulate. We previously demonstrated that CK2α’ is progressively upregulated in the striatum of HD mice and that its levels correlate positively with mHTT aggregation, with the strongest association starting at 12 months of age (25). Therefore, our findings suggest that CK2 inhibition exerts its greatest therapeutic benefit at later disease stages, when CK2α’ upregulation and mHTT aggregation are more pronounced. This also suggests that up-regulation of CK2α’ may not be directly related to the onset of mHTT aggregation but rather with the accumulation of aggregates over time.

The impact mediated by CK2α’ in mHtt aggregation can be due to a variety of different mechanisms. CK2α’ controls the regulation of Heat shock factor 1 (HSF1), the master regulator of protein homeostasis (24,71,73). Increased CK2α’ in HD causes the inactivation and proteasomal degradation of HSF1 through inhibitory phosphorylation and resultant ubiquitination (24,73). It has been shown that depletion of HSF1 in HD contributes to the depletion of chaperones expression and increases protein aggregation (74). Inhibition of CK2 may restore protein homeostasis by restoring HSF1 and chaperones expression. CK2 also participates in the regulation of autophagy by phosphorylating autophagy regulators such as ULK1 and TFEBI (75,76) and therefore, inhibition of CK2 can result in increased Htt clearance. It has also been reported that CK2 phosphorylates Htt directly through phosphorylation of N17, altering Htt aggregation (70,72,77), although the exact role of CK2α’ in this process is not known. It is therefore possible that CK2 inhibition impacts Htt aggregation through a myriad of mechanisms that can simultaneously operate in the cell by directly modulating conformational states in Htt less prone to aggregation, by restoring overall folding capacity and/or increasing protein clearance. Future studies will be essential to uncover the mechanisms by which CK2 inhibition mediates its beneficial effects in HD.

mHtt aggregation has been shown to be accompanied by hypoexcitability of MSNs and decrease excitatory synapse density (78–81). Importantly, we found that decreased neuronal mHtt aggregation in CX-4945-treated HD mice, especially at symptomatic stages, was accompanied by an increase in DARPP-32 and cFos expression and increased cortico-striatal synapse density (PSD-95/VGLUT1). Similar results have been found when genetically deleting CK2α’ in zQ175 mice (25). However, whether these effects are mediated by a reduction in mHtt aggregation or by the direct action of CK2 inhibition is unclear. CK2 directly phosphorylates several proteins involved in synapse stability and synaptic plasticity including cFos, PSD-95, and CAMKII which in turn regulate glutamatergic receptor trafficking and neuronal signaling (82–85). The expression of PSD-95 and VGLUT1 are also impacted by the levels of CK2α’ since HD mice lacking one allele of CK2α’ also presented enhanced levels of PSD-95 and VGLUT1 (25). Therefore, it is possible that the effects mediated by CX-4945 in cFos expression and cortico-striatal synapse density are due to a direct impact in the phosphorylation of these substrates. We also found that the depletion of cFos expression in HD mice at 6 months did not necessarily correlate with an alteration in cortico-striatal synapse density, but it was rescued in CX-4945 treated mice. This is possibly due to alterations in other types of striatal synapses such as thalamo-striatal synapses, which are also depleted in HD mice and at earlier stages than cortico-striatal synapses, and rescued by CK2α’ haploinsufficiency (25,78).

CK2 has also been shown to be a key regulator of neuroinflammation through the phosphorylation of NF-kB (70,86). Treatment with CX-4945 in symptomatic mice reduced the expression of multiple inflammatory cytokines and normalized microglial morphology by restoring the number of microglial branches. Given that the upregulation of CK2α’ in HD mice is largely restricted to neurons, with little to no detectable expression in glial cells (24), it is reasonable to conclude that the observed improvements in striatal inflammatory state and microglial morphology may arise from neuronal modulation of CK2α’ activity following CX-4945 treatment. This suggests that the changes observed in glial state likely reflect a secondary, non–cell autonomous response to neuronal dysfunction driven by mHtt and its associated CK2α’ elevation.

A reduction of neuronal CK2 activity and the accompanying decrease in inflammatory cytokines can also have a stabilizing effect on astrocytic homeostatic functions (87,88). The reduction of homeostatic function of HD astrocytes has been established in previous studies (13,16,89). These functions include a decrease in calcium signaling, glutamate uptake, and phagocytosis. Under pathological conditions, elevated neuronal CK2 activity promotes NF-κB–dependent cytokine release (e.g., TNFα, IL-1β, IL-6), which act on astrocytes to induce reactive phenotypes characterized by increased GFAP and S100B expression (90–92). We previously found that CK2α’ knockdown in zQ175 mice reduced expression of several cytokines including TNFα, IL-1β and IL-6 and reversed transcriptional dysregulation of several astrocyte genes associated with calcium signaling, glutamate uptake and phagocytosis (25). Contrary to what it was expected, we found that CX-4945 treatment increased the number of S100B^+^ astrocytes, which can be interpreted as an increase in reactive astrogliosis. However, increasing evidence indicates that moderate elevations of S100B can also exert neuroprotective effects (58,59,63). We found that CX-4945 treatment enhanced the accumulation of cytoplasmic mHTT aggregates in S100B^+^ astrocytes, and that this was potentially associated with enhanced phagocytic activity by these astrocytes, ultimately reducing mHtt burden and contributing to increase neuronal health. In support to our findings, studies in other neurodegenerative diseases have also reported that depletion of CK2-dependent targets also increased number of reactive astrocytes but restored phagocytosis and depleted reactive microgliosis (93–96). Additionally, S100B contributes to Ca^²⁺^ buffering and redox homeostasis within astrocytes, supporting their homeostatic functions during stress (97,98). Therefore, an increase in S100B may reflect a compensatory or protective astrocytic response, aimed at preserving neuronal integrity and maintaining homeostasis. To this extent, we found that CX-4945 increased Ca^²⁺^ transients and amplitude of events in the branches of astrocytes of HD-treated mice. Taken together, these data suggest that CK2 inhibition restores homeostatic function of astrocytes. CK2 upregulation in HD appears to be largely neuronal (25), suggesting that the beneficial effects of CX-4945 on microglia and astrocyte homeostasis are mediated indirectly, via neuron-glia signaling. However, to fully understand the beneficial effects exerted by CX-4945 in HD mice, it will be important to parse the role of CK2α’ in cell-autonomous vs non-cell-autonomous impacts on neurons, astrocytes, and microglia.

Importantly, the beneficial effects mediated by CX-4945 in the pathology of HD mice were translated to improved motor performance, ameliorating deficits in beam walk and foot fault tests. Although the most profound effects in the amelioration of striatal pathology by CX-4945 were found in symptomatic groups, we found improved motor behavior in both prodromal and symptomatic treated mice, emphasizing the relevant functions that CK2α’ play in HD pathophysiology in both onset and disease progression.

Overall, our findings demonstrate that CX-4945 treatment ameliorates key pathological hallmarks of HD, including mHtt aggregation, neuronal dysfunction, and neuroinflammation. Notably, the therapeutic effects were more pronounced when treatment was initiated at late symptomatic stages, suggesting that CX-4945 may remain effective even after disease onset. This is extremely encouraging given that HD patients are typically diagnosed after the appearance of clinical symptoms, implying that CX-4945 could offer meaningful therapeutic benefit within a clinically relevant window. Moreover, CX-4945 has already demonstrated safety and tolerability in humans, having been used in phase 1 and 2 clinical trials for other diseases (99–101). While additional studies are required to establish its full efficacy and mechanism of action in HD, these results provide a strong foundation for advancing CK2 inhibition as a promising therapeutic strategy that could be rapidly translated to the clinic.

---

**Supplementary Table 1:**
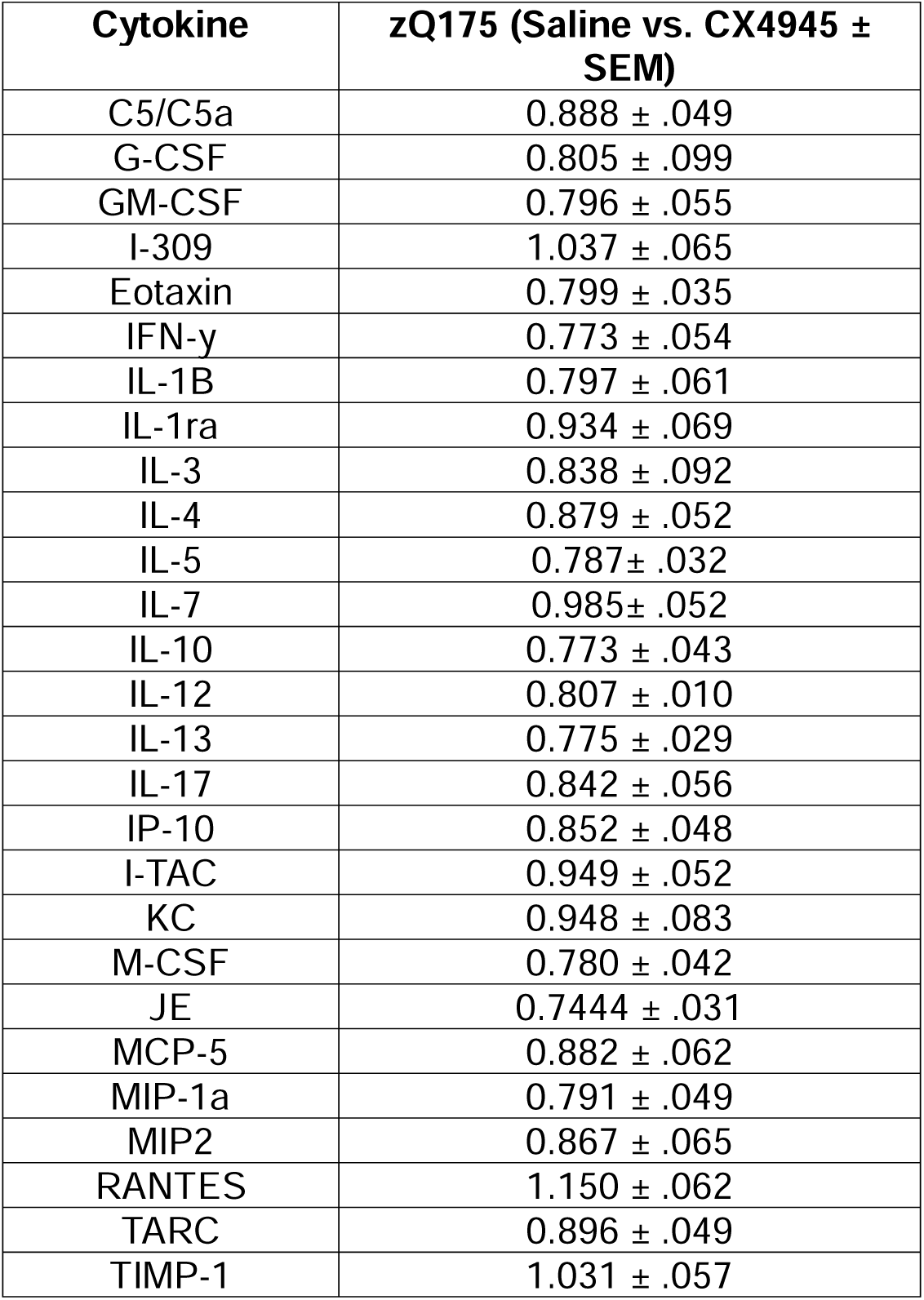
Cytokines from the cytokine proteome profile that did not show significant differences between treatments.

## Author Contributions

RGP obtained funding for this study. RGP and RJP designed the experiments. RJP performed oral gavage treatment, behavioral studies and Calcium imaging. RJP, MH, NBR and MAS conducted immunofluorescence experiments. RJP, MH, NBR and MAS prepared and analyzed the data. RJP and RGP interpreted the data and wrote the first draft of the manuscript. All other authors edited subsequent versions and approved the final version of the manuscript.

## Acknowledgements

We thank Dr. Ezequiel Marron at the University of Minnesota Viral Core for his assistance in viral production, Dr. Mark Sanders and Dr. Jason Mitchell at the University of Minnesota University Imaging Center for help accessing imaging equipment and providing technical support and Dr. Erin Lind at the Mouse Behavior Core for her assistance in accessing mouse behavior equipment. We thank Drs. Paulo Kofuji, Alfonso Araque and Julianna Goenaga for their help with 2P-imaging and analyses.

